# Rapid phonetic learning of Mandarin tones in adults: Daily behavioral improvement and brain activity changes

**DOI:** 10.1101/2025.10.06.680317

**Authors:** Xueqiao Li, Hyeonjeong Jeong, Piia Astikainen

## Abstract

Although adults learn foreign languages more slowly than children, behavioral improvements can still emerge rapidly with training. Previous phonetic learning studies have mostly focused on pre- and post-training comparisons, leaving daily learning trajectories largely unexplored. Here, in Finnish speakers naïve to tone languages, we tracked day-to-day changes in Mandarin tone perception during a short training program (1 h/day for 4 days), complemented by pre- and post-training behavioral and event-related potential (ERP) measures. Each learning session comprised exposure to Mandarin tones in continuous and isolated speech with cues, change detection and identification with feedback, and a listen-and-repeat exercise. To assess the role of co-presence, participants learned either in pairs (n = 22) or individually (n = 20). We found that discrimination and categorization speed, as well as discrimination accuracy, improved from pre-to posttest, with performance also increasing across daily training sessions. Paired learners showed higher sensitivity (d′) to tone changes on the first day than individual learners, consistent with co-presence–driven attentional facilitation. At the neural level, P3a amplitude to tone changes during passive listening increased after training, reflecting enhanced automatic orienting to novel sounds at whole group level. These findings demonstrate that short-term training induces rapid behavioral gains and selective neural plasticity in adult phonetic learning, with early co-presence effects and transfer to novel speech sounds.

## INTRODUCTION

A fundamental challenge in adult second language (L2) acquisition is the accurate perception of phonetic features that are absent in the learner’s native language. While infants are initially sensitive to a wide range of speech contrasts, this universal perceptual sensitivity becomes attuned to the features of the ambient language through early exposure, leading to reduced discrimination of non-native contrasts over time (e.g., Cheour et al., 1998; Kuhl et al., 1992; Polka & Werker, 1994; Werker & Tees, 1984). This perceptual narrowing limits the ability to perceive unfamiliar phonetic contrasts later in life, making the acquisition of novel phonetic categories more demanding for adult L2 learners. A notable example is the acquisition of lexical tones, which convey word meaning in tonal languages such as Mandarin but are absent in non-tonal languages like English and Finnish. Learners from non-tonal language backgrounds often show persistent difficulty in perceiving, categorizing, and producing tonal distinctions (Hao, 2012; Hallé et al., 2004; Pelzl et al., 2019, 2021).

Despite persistent challenges in adult phonetic learning, we know from experience, that adults can learn new languages. There is also growing research evidence that targeted training protocols (e.g., Baills et al., 2019; Callan et al., 2003; Tremblay et al., 1998) and extended passive exposure (e.g., Kurkela et al., 2019; Li et al., 2025) can induce plastic changes in the cortical processing supporting change detection of foreign phonetic contrasts. However, behavioral identification, discrimination and change detection accuracy and concurrent brain activity changes related to phonetic learning are not well understood. Some training studies have shown both behavioral and brain activity changes (e.g., Callan et al., 2003; Cheng et al., 2019; Tamminen et al., 2015), while sometimes brain activity changes have been reported without behavioral gains (e.g., Kurkela et al., 2019; Tremblay et al., 2014). In addition, many studies have investigated only either behavioral (e.g., Huensch & Tremblay, 2015; Melnik & Peperkamp, 2021; Olson, 2019) or brain activity changes (Li et al., 2025) related to phonetic learning of foreign speech sounds. Understanding the temporal dynamics and neural mechanisms of phonetic learning and their relationship with behavioral improvements is critical for theories of auditory processing, second-language acquisition, and experience-dependent brain plasticity.

The learning-related changes are often tracked using electrophysiological (EEG) measures such as event-related potentials (ERPs), which provide high temporal resolution for examining auditory learning dynamics. A widely used method is measuring speech sound change detection by using so called oddball paradigm, in which frequent standard stimuli are occasionally replaced by infrequent deviant stimuli. Within this framework, different ERP components are associated with distinct stages of auditory and cognitive processing. The mismatch negativity (MMN), occurring approximately 150–250 ms after stimulus onset, reflects automatic detection of acoustic change during passive listening and is typically maximal at fronto-central scalp sites (Näätänen et al., 1978; Näätänen et al., 2007). The subsequent P3a component, peaking around 250–500 ms, is associated with involuntary orienting of attention toward salient or unexpected sounds and is often observed as a fronto-central positivity (Polich, 2007). In tasks requiring active detection of deviant sounds, ERPs such as the N2b and P3b emerge. The N2b, typically observed around 200–350 ms with a fronto-central distribution, reflects controlled detection of stimulus deviance; while the P3b, peaking around 300–600 ms with a centro-parietal distribution, is linked to the updating of working memory (Folstein & Van Petten, 2008; Kok, 2001; Polich, 2007).

In the context of foreign language phonetic learning, ERP components recorded in the oddball condition are sensitive to learning resulting from exposure and training in adults. For instance, passive exposure to Mandarin lexical tones over multiple days has been shown to enhance the P3a response in native Finnish speakers, suggesting increased involuntary attentional orienting toward novel tonal contrasts (Kurkela et al., 2019). However, findings on the MMN remain mixed: while some studies report greater MMN as a result of improved deviance detection (Näätänen et al., 1993; Kraus et al., 1995; Tamminen et al., 2015; Tremblay et al., 1997), others report decreased MMN responses (Lu et al., 2015; Li et al., 2025) or no observable changes in MMN amplitudes (Kaan et al., 2008; Kurkela et al., 2019) after training or exposure. When learners are actively engaged in detecting tonal deviations, later components such as the P3b have been shown to increase following training, reflecting the consolidation of newly acquired phonetic distinctions (Alain et al., 2010; Giroud et al., 2017; Kurkela et al., 2019), though decreased responses have also been observed (Ben-David et al., 2011). Learning-related modulations of the N2b also appear to depend on task demands and training duration. For example, identification training with sufficient intensity has been associated with N2b enhancement (Alain et al., 2010), whereas shorter (e.g., one hour) training or passive exposure alone often fails to produce N2b amplitude changes (Ben-David et al., 2011; Giroud et al., 2017; Kurkela et al., 2019). However, few ERP-studies have utilized both passive and active change detection, leaving it unclear whether learning related changes are similar for both (see however Kurkela et al., 2019 for effects of passive exposure). Furthermore, most previous studies have examined learning outcomes using only pre- and post-training comparisons (e.g., Callan et al., 2003; Cheng et al., 2019; Huensch & Tremblay, 2015; Melnik & Peperkamp, 2021; Olson, 2019). To the best of our knowledge, only one recent study has attempted to trace the phonetic learning dynamics *during* training, but this study was based on passive listening (Li et al., 2025) and did not include behavioral assessment of phonetic learning.

To address these gaps in the literature, the present study examined how a four-day training of Mandarin lexical tones shapes oddball ERP responses in passive and active listening, as well as behavioral outcomes in discrimination and identification, in adult Finnish speakers with no prior experience of tonal languages. The discrimination task measured perceptual sensitivity to acoustic differences between tonal stimuli, providing insight into learners’ ability to detect subtle pitch variations. The categorization task required participants to assign each tone to a predefined phonological category (e.g., rising or falling), thereby probing whether learners were able to form stable, abstract representations of foreign tone contrasts. A previous study has reported that after training, learners tend to show more accurate responses and faster reaction times in tone categorization (Zhang et al., 2024), reflecting the successful formation of new phonological categories. Similarly, training has been shown to improve the discrimination of tone pairs (Chandrasekaran et al., 2010). However, little is known about the time course of learning for these two processes. The ability to discriminate speech sounds does not necessarily guarantee successful categorization, as the latter requires mapping acoustic variation onto linguistically meaningful categories (for categorical perception, see Goldstone & Hendrickson, 2010). By combining these measures, we can investigate whether perceptual sensitivity and phonological categorization develop in parallel during short-term training, or whether early gains in detection precede the emergence of robust categorical representations. This question has direct implications for models of adult speech learning and lexical processing.

To capture the dynamics of learning and its different mechanisms, behavioral responses during the training phase were recorded for identification and change detection, allowing us to trace day-by-day changes in perceptual sensitivity. The identification task was virtually the same as the categorization task in the pre- and post-tests, but during learning sessions only clearly rising and falling tones were used in several different syllables to train the participants. The change detection task resembled the active oddball condition in the pre- and post-tests, except that it employed the consonant-vowel syllable /fa/.

Language learning often occurs in a social context, yet relatively little is known about adult social learning of phonetic features. Interacting with peers can enhance motivation, attention, and memory, and recent work in educational neuroscience highlights the role of joint attention and social engagement in shaping learning outcomes (Verga & Kotz, 2013; Li & Jeong, 2020). While vocabulary learning in social contexts has been studied extensively, with clear advantages in interactive conditions (Verga et al., 2015; Verga & Kotz, 2017, 2019), it remains unclear whether similar benefits extend to phonetic learning. To address this question, the present study compared learning outcomes between adults who trained alone and those who trained in pairs.

In general, we hypothesize that a 4-hour training to Mandarin lexical tones will lead to measurable changes at both behavioral and neural levels. Specifically, we expect improvements in categorization and discrimination accuracy, as well as faster response times in the post-compared with pre-training measurement. We further predict that discrimination gains would emerge earlier than identification improvements, reflecting a potential sequential development of perceptual and phonological representations. We also expect enhanced ERP responses following training. Based on previous findings, we anticipate observing a larger amplitude of P3a in passive listening and N2b and P3b in active listening in the post-compared to the pre-training measurement. We also hypothesize that pair-based learning will facilitate phonetic acquisition, resulting in either earlier or more robust learning effects, potentially through mechanisms such as increased attention, motivation, and engagement.

## METHODS

### Participants

The participants were recruited through announcements on social media and through the University of Jyväskylä’s email lists. Written informed consent was obtained from all participants before their participation. The experiment was undertaken in accordance with the Declaration of Helsinki. The ethical committee of the University of Jyväskylä approved the research protocol.

Inclusion criteria for the study were right-handedness and age of 18-40 years. Exclusion criteria were exposure to tonal languages (no more than two weeks of passive exposure, for example, during a holiday trip), language impairment, such as reading, writing or speech impairment, or sensory impairment (corrected vision allowed), learning disability, attention deficit disorder, neurological disease or disability, motor disorder, psychotic disorder, autism spectrum disorder, social anxiety disorder, or recent drug use or abuse, or medication affecting the central nervous system.

The study involved 42 Finnish native speakers divided into two groups: Individual (*n* = 20; 14 females and 6 males; mean age = 24.65 years, *SD* = 5.18) and Pair (*n* = 22; 14 females and 8 males; mean age = 24.96 years, *SD* = 4.84). The sample size is consistent with prior ERP studies on phonetic learning of Mandarin tones using the oddball paradigm (e.g., Kurkela et al., 2019; Li et al., 2025; Shen & Froud, 2019) and is comparable to that used in studies examining social co-presence effects in word learning (Verga & Kotz, 2017, 2019).

There were no significant differences between the groups in demographic variables (age, gender, education level) or in psychological measures, including scores on the Beck Depression Inventory-II (BDI-II), Beck Anxiety Inventory (BAI), State-Trait Anxiety Inventory-Y2 (STAI-Y2), and Social Provision Scale (SPS). Additionally, participants reported similar levels of prior experience with musical instruments, including singing.

In the final analysis, for the behavioral discrimination task in the pre- and post-tests, only 19 participants in the individual group and 22 in the pair group were included due to missing data. For the ERP data, 20 participants from the individual group and 21 from the pair group were included in the passive change detection task, and 20 from each group were included in the active change detection task after excluding data with excessive artifacts such as eye movements.

### Stimuli

Two sets of auditory stimuli were used in this study: those applied during the learning phase and those presented in the pre- and post-tests.

For the learning phase, natural speech stimuli were used. These included both continuous and isolated speech materials. Continuous speech consisted of short poems, while isolated speech comprised consonant–vowel syllables produced with three distinct Mandarin Chinese tones: flat (lexical tone 1), rising (lexical tone 2), and falling (lexical tone 4). All speech materials were recorded in both audio and video formats by a female native speaker of Mandarin Chinese. A complete list of the stimuli used during the learning phase is provided in the supplementary materials.

The auditory stimuli used in the pre- and post-tests consisted of lexical tones based on the vowel /a/. These tonal stimuli were prepared by recording a female native Mandarin Chinese speaker articulating the vowel /a/ with either a rising (Mandarin Chinese lexical tone 2) or a falling (Mandarin Chinese lexical tone 4) pitch contour. Recordings were made at a sampling rate of 44.1 kHz. The sounds were digitally edited using SoundForge software (SoundForge 9, Sony Corporation, Japan) to standardize their duration to 200 ms. To isolate pitch contour as the primary variable while keeping all other acoustic features constant, pitch tier transfer was performed using Praat software (version 5.4.06; University of Amsterdam). This procedure generated a rising tone and a falling tone that were acoustically identical except for their fundamental frequency (F0) contours. These two endpoint tones were then used to create a continuum of 11 lexical tone tokens separated by 10 equal interval steps. The detailed procedure for stimulus generation has been reported previously (Kurkela et al., 2019; Xi et al., 2010).

From this continuum, six tone tokens (Tokens a1, a3, a5, a7, a9, and a11) were selected for use in the behavioral discrimination and categorization tasks. These tokens represented a graded progression from a distinctly rising tone (Token a1) to a clearly falling tone (Token a11), with intermediate steps reflecting gradual pitch transitions.

For the EEG measurements, a subset of three stimuli was selected from this continuum. The standard stimulus (Token a11) was a distinctly falling tone, with an F0 decreasing from 326.9 Hz to 162.1 Hz. The small deviant (Token a7) featured a moderately falling contour from 274.4 Hz to 210.2 Hz, and the large deviant (Token a3) was a rising tone, with F0 increasing from 226.2 Hz to 268.4 Hz. Sound levels were calibrated using a Brüel & Kjær sound level meter (2250 series), with direct measurements taken through an insert earphone connected to an ear simulator (Type 4157-X, ID No. 26767). The A-weighted equivalent continuous sound level (LAeq) was 73.2 dB, and the A-weighted fast maximum level (LAFmax) was 78.3 dB.

### Procedure

The experiment employed a pre–post design, with four consecutive days of learning in between. All participants completed the pre- and post-tests individually, regardless of their group assignment. These tests included both behavioral assessments and EEG recordings. During the learning phase, participants attended one hour of training per day for four consecutive days and completed the training individually or in pairs, depending on their group assignment. The details of experimental design are shown in Figure 1.

**Figure 1.**
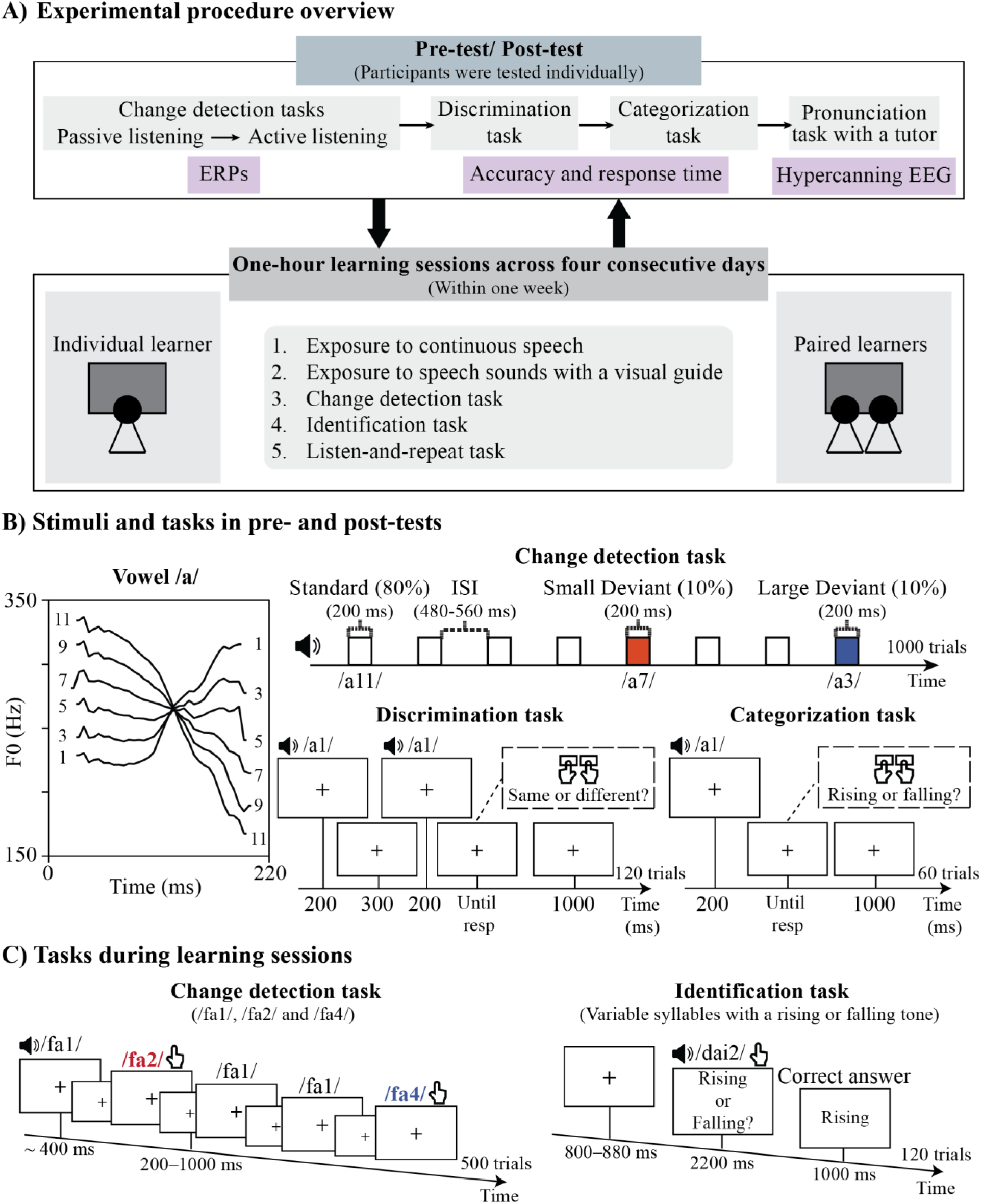
Illustration of the experimental design and example trials from tasks measuring learning effects. A) Overview of the experimental procedure. All participants completed the pre- and post-tests individually, with four consecutive days of learning in between. Hyperscanning EEG results during the pronunciation task in the pre- and post-tests are not reported here. B) Fundamental frequency (F0) of the stimuli and example trials for each testing task. In the change detection task, tokens a3, a7, and a11 were used. In the discrimination and categorization tasks, tokens a1, a3, a5, a7, a9, and a11 were used. *resp* = response. C) Example trials from the change detection and identification tasks during the learning sessions. The numbers next to the syllables represent lexical tones in Mandarin Chinese: 1 = flat, 2 = rising, 4 = falling. In the identification task, questions and correct answers were presented in Finnish. For illustration purposes, the examples shown in the figure are translated into English.

### Experimental procedure in learning sessions

The same procedure was followed on each of the four training days. Individual learners and paired learners used the same set of tasks and materials; however, participants in the paired group completed the tasks at a shared computer, using a single keyboard with response keys positioned at opposite ends, while viewing the same monitor (27” with a desktop resolution of 1920 × 1080).

Each learning session began with an exposure phase featuring continuous natural speech. Participants watched videos lasting approximately 10 minutes, during which several Chinese poems were recited. A different set of poems was presented each day to ensure novelty and maintain engagement. After the continuous speech exposure, participants were presented with 60 isolated consonant–vowel syllables with three different tones (i.e., flat, rising, falling), one by one in a random order, and each accompanied by a visual cue indicating its corresponding tone category.

After the exposure phases, participants completed three interactive tasks. The first was a change detection task, in which the syllable /fa/ was presented repeatedly using an auditory oddball paradigm. The standard stimulus was a flat tone (/fa1/), presented in 80% of the trials, while rising and falling tones (/fa2/ and /fa4/, respectively) served as the deviant stimuli (each presented in 10% of the trials). Participants were instructed to press a response key whenever they heard a tone that differed from the standard. A total of 500 stimuli were presented in pseudorandom order that deviant tones were separated by at least two standard trials. Participants received feedback on their overall accuracy at the end of the task.

The second task was a tone identification task, in which participants heard 60 consonant–vowel syllables produced with either a rising or falling tone and categorized each by pressing the corresponding button, resulting in 120 trials. The mapping of response keys was counterbalanced across participants. Feedback was provided on each trial, as well as in summary form at the end of the task, to support learning.

The final task was a listen-and-repeat task. Participants watched short video clips of a native speaker articulating consonant–vowel syllables and were instructed to repeat each syllable aloud, either alone or together with the partner. Each syllable was produced with a flat tone, a rising tone, and a falling tone. In total, 180 stimuli were presented in a random order.

Auditory stimuli during these tasks were delivered via headphones, and the presentation and timing of the stimuli were controlled using PsychoPy (Version 2022.2.2; Peirce et al., 2019).

In addition to the tasks, questionnaires assessing participants’ alertness, motivation, self-evaluation of performance, and perceived pleasantness of the learning experience were measured. A detailed description of these measures, along with the corresponding analyses and results, is provided in the supplementary materials.

### Experimental procedure in pre- and post-tests

#### Change detection tasks during EEG recordings

During both the pre- and post-test sessions, EEG measurements were conducted prior to the behavioral tasks. Two experimental conditions were applied in a fixed order, with the passive condition preceding the active condition. In the passive condition, participants were instructed to watch a muted video with Chinese subtitles and to focus solely on the video while ignoring any sounds they heard. In the active condition, participants were instructed to detect deviant tones by pressing a response key as quickly and accurately as possible.

In both conditions, auditory stimuli were presented via earphones in an oddball paradigm. The stimulus set consisted of one standard tone (Token a11, presented on 80% of trials), one small deviant tone (Token a7, presented on 10% of trials), and one large deviant tone (Token a3, presented on 10% of trials). A total of 1000 stimuli were delivered in pseudorandom order, with the constraint that each deviant was preceded by at least two standard stimuli. The interstimulus interval (ISI) varied randomly between 480 and 560 ms (offset-to-onset).

The presentation of auditory stimuli was controlled using Presentation software (Version 23.0 Build 10.27.21, Neurobehavioral Systems, Inc.). Throughout both conditions, participants were seated comfortably in a well-lit room. Continuous EEG was recorded using the NeurOne amplifiers (Bittium Biosignals Ltd., Kuopio, Finland) with 64-channel caps (Easycap GmbH, Wörthsee, Germany). The sampling rate during recording was 1000 Hz and the data were filtered online with a 250 Hz high cut-off. EEG was referenced online to FCz.

#### Tone discrimination and categorization tasks

Two behavioral tasks were conducted following the EEG measurements: a tone discrimination task and a tone categorization task. All participants completed the discrimination task first, followed by the categorization task. The discrimination task was given first to make sure participants judged the sounds based on perception, without being influenced by category labels. Auditory stimuli were delivered through over-ear headphones, with presentation and timing controlled by PsychoPy (v2021.2.3; Peirce et al., 2019).

The tone discrimination task employed an AX paradigm, where participants judged whether two sequentially presented /a/ syllables had the same or different tones. Participants were instructed to respond as quickly and accurately as possible by pressing either the “F” or “J” key, with key assignments counterbalanced across participants. A total of 120 trials were presented, consisting of 60 same-sound pairs and 60 different-sound pairs. Same pairs included 10 repetitions of each token pairing (i.e., Token a1–Token a1, Token a3–Token a3, Token a5– Token a5, Token a7–Token a7, Token a9–Token a9, and Token a11–Token a11), while different pairs included 12 repetitions of each token pairing (i.e., Token 1–Token 3, Token 3– Token 5, Token 5–Token 7, Token 7–Token 9, and Token 9–Token 11). Each trial consisted of a sound pair separated by a 300 ms inter-stimulus interval. A new trial began 1000 ms after the participant’s response. The presentation order of the sound pairs was randomized across trials. A short break was provided after the first 60 trials to minimize fatigue. Both accuracy and response time were recorded for each trial.

The tone categorization task began with a familiarization phase, during which participants listened to labeled examples of the most rising and most falling tones (Tokens a1 and a11). After the familiarization, /a/ syllable with different tones was presented one at a time.

Participants were instructed to categorize each tone as either rising or falling by pressing the “F” or “J” key as quickly and accurately as possible. Key assignments were counterbalanced across participants. Each of the six tokens from the tone continuum (Tokens a1, a3, a5, a7, a9, and a11) was presented 10 times, resulting in a total of 60 trials. The order of the trials was randomized. A fixed inter-trial interval of 1000 ms followed participants’ each response. Response times and categorical decisions were recorded for analysis.

### Data analysis

#### Behavioral data analysis

Behavioral responses, including accuracy (ACC) and response time (RT), were extracted from tasks completed during the learning process as well as the pre- and post-tests. Only responses with RTs longer than 200 ms were considered valid. For RT analyses, only valid and correct responses were included.

For the change detection task during the learning process and the active condition in change detection task during the pre- and post-tests, perceptual sensitivity (*d*′, d prime) was calculated (*d*′=z(hit rate)−z(false alarm rate)). Valid responses to deviants were marked as hits, while no responses to deviants were classified as misses. Responses to standards were marked as false alarms, and no responses to standards were considered correct rejections. In addition, responses with RTs shorter than or equal to 200 ms were treated as invalid and reclassified as misses for deviant trials and false alarms for standard trials. For the identification task during the learning process, response accuracy and response times were analyzed.

For the discrimination task during the pre- and post-tests, participants judged whether pairs of sounds were the same or different. Each trial was classified as either a same-pair or different-pair trial, and this categorization was used in the statistical analysis.

For the categorization task during the pre- and post-tests, participants judged whether the tone of each sound was rising or falling. In the analysis, the tone tokens were grouped into these categories based on their physical features: rising (tokens 1 and 3) and falling (tokens 9 and 11). Participant responses were analyzed based on the accuracy of assigning each token to its correct tone category (rising or falling).

#### EEG data analysis

EEG preprocessing was performed using MNE-Python (version 3.11.5; Gramfort et al., 2013). Data were filtered with a bandpass of 0.1–30 Hz. A notch filter of 50 Hz was applied to reduce electrical noise. Bad channels were manually identified via visual inspection and interpolated using a spherical spline model (Perrin et al., 1989). The data were then re-referenced to the average of all EEG channels. Independent Component Analysis (ICA) was conducted using the FastICA algorithm (Hyvärinen, 1999) to detect and remove eye movement artifacts from the continuous data. Epochs were extracted for each deviant type and for the standard stimuli that immediately preceded the corresponding deviant stimuli, from –100 ms to 600 ms relative to stimulus onset. Baseline correction was applied from –100 to 0 ms. Trials with peak-to-peak amplitudes exceeding 75 μV or flat activity below 0.5 μV were excluded. Epochs were averaged separately for each stimulus type (small and large), deviant type (deviant and standard), session (pre and post), and participant. Only data of the participants with more than 50 accepted epochs per condition were included in the analysis.

MMN and P3a were analyzed in the passive oddball condition, and N2b and P3b in the active oddball condition. Time windows for statistical analysis were guided by prior reviews (Folstein & Van Petten, 2008; Näätänen et al., 2007; Polich, 2007) and defined from the grand-averaged ERPs: 140–190 ms for MMN, 230–290 ms for P3a, 190–230 ms for N2b, and 320–390 ms for P3b. Mean amplitudes were calculated at electrode sites selected based on visual inspection of the topographical distributions, consistent with earlier work (Kurkela et al., 2019): F3, Fz, and F4 for MMN and P3a; FC1, FC2, C3, Fz, and C4 for N2b; and CP1, CPz, and CP2 for P3b.

#### Statistical analysis

Statistical analyses of behavioral data were conducted in RStudio (version 2024.12.1+563) using R (version 4.4.3). The lme4 package (version 1.1-36) was used to fit both linear mixed-effects models (LMMs) and generalized linear mixed-effects models (GLMMs).

For RTs, LMMs were applied when the data were not strongly skewed (skewness ≤ 1). If the data were positively skewed (skewness > 1), a GLMM with a Gamma distribution and log link function was used. For accuracy data, which were coded as binary outcomes (correct = 1, incorrect = 0), a binomial GLMM was applied. For *d*′ scores, LMMs were used. To address model singularity, by-item random slopes and intercepts were removed in a stepwise fashion. The optimizer “bobyqa” was used to resolve convergence issues when necessary (Brauer & Curtin, 2018). Details of model structures are reported in the Results section.

For both GLMMs and LMMs, the significance of fixed effects and their interactions was assessed using Type III Wald χ² tests, implemented with the *Anova* function from the *car* package (version 3.1-3). To complement frequentist inference, Bayesian model comparisons were conducted for group- and day/session–related main and interaction effects to evaluate whether groups differed in their learning performance and whether there was a general learning effect across sessions. These comparisons were performed by contrasting full models with reduced models in a hierarchical order. Models were refitted using maximum likelihood (ML) estimation, and Bayes factors (*BF*_10_) were approximated from Bayesian Information Criterion (BIC) differences (Wagenmakers, 2007). Following conventional thresholds, *BF*_10_ > 3 was considered strong evidence, 1 < *BF*_10_ < 3 weak evidence, and *BF*_10_ < 1 evidence against the effect.

In practice, post hoc comparisons were only conducted when higher-level tests were supported (*p* < .05 and *BF*_10_ > 1). If the frequentist test was non-significant (*p* ≥ .05) but the Bayes factor suggested weak evidence (1 < *BF*_10_ < 3), the result was treated as inconclusive, whereas strong Bayesian evidence (*BF*_10_ > 3) was taken to support further post hoc testing.

In the presence of significant three-way interactions, interaction analyses were conducted using the joint_tests function from the emmeans package (version 1.10.7) to examine two-way interactions at each level of the third variable.

Estimated marginal means were used for post hoc comparisons. For GLMMs, pairwise comparisons were conducted using Wald *z* tests; for LMMs, Satterthwaite-approximated t-tests were used. Degrees of freedom and *p*-values for LMMs were estimated using Satterthwaite’s approximation, as implemented in the lmerTest package (version 3.1-3). Bonferroni correction was applied to adjust for multiple comparisons. For significant results, parameter estimates (β), standard errors (*SE*), test statistics, *p*-values, and 95% confidence intervals (*CI*s) are reported.

Statistical analyses of ERP data were conducted using JASP (Version 0.18.2; JASP Team, 2023). Mean amplitude values were analyzed separately for each ERP component using a four-way repeated-measures ANOVA, with stimulus type (deviant vs. standard), deviant type (small vs. large), and session (pre vs. post) as within-subject factors, and group (individual vs. pair) as a between-subject factor. Partial eta squared (partial η²) was reported as an index of effect size for *F*-tests. Pairwise comparisons were conducted to further examine significant interaction effects, using estimated marginal means to compare main effects or simple main effects, with Bonferroni adjustment applied to the confidence intervals. For significant post hoc comparisons, we report *p*-values, 95% *CI*s, and Cohen’s *d* as the effect size.

All reported mean values in the Results section are estimated marginal means with standard errors. For all statistical tests on behavioral data, significance was defined as *p* < .05.

## RESULTS

### Results from tests during learning sessions

The results indicating group differences and learning effects are summarized and depicted in Figure 2.

**Figure 2.**
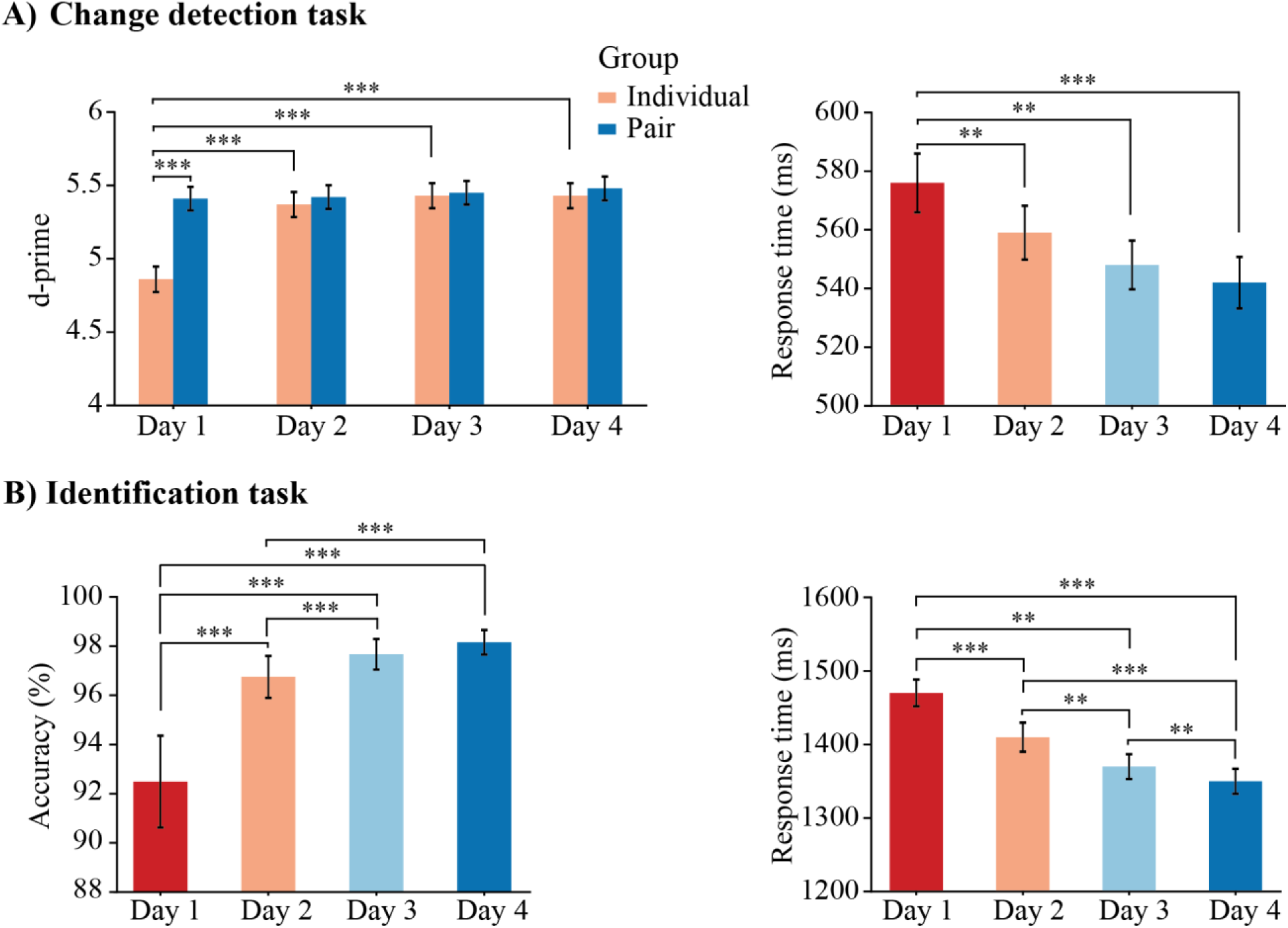
Results during the learning process. A) Results from the change detection task. Post hoc results for the interaction effect between group and day in d-prime (left), as well as pairwise comparisons examining the main effect of day on response time (right), are illustrated. B) Results from the identification task. Results investigating the main effect of day in accuracy (left) and response time (right) are illustrated. All bar charts display the estimated marginal means (EMMs), with error bars representing the standard error (*SE*) of the EMMs. ** = p < .01, *** = p < .001.

### Change detection task

#### Accuracy

For accuracy, d-prime (*d′*) was applied and analyzed using a linear mixed-effects model (LMM) with a Type III Wald χ² test. The model included group (individual vs. pair) and day (1, 2, 3, 4) as fixed effects. Participant was included as a random effect with a random intercept. The analysis revealed a significant main effect of group (Wald χ²(1) = 21.26, *p* < .001), a significant main effect of day (Wald χ²(3) = 52.97, *p* < .001), and a significant group × day interaction (Wald χ²(3) = 24.26, *p* < .001). Bayesian model comparisons strongly supported the inclusion of the group × day interaction (*BF*_10_ = 51.654) and the main effect of day (*BF*_10_ = 63.447), whereas the main effect of group was not supported (*BF*_10_ = 0.408).

Pairwise comparisons within each group revealed a significant increase in sensitivity (*d′*) across days for the individual group. Significant differences were observed between day 1 (*M* = 4.86, *SE* = 0.087) and day 2 (*M* = 5.37, *SE* = 0.085; β = 0.507, *SE* = 0.092, *t*(120) = 5.49, *p* < .001, 95% *CI* [-0.754, -0.259]), day 1 and day 3 (*M* = 5.43, *SE* = 0.085; β = 0.572, *SE* = 0.092, *t*(120) = 6.20, *p* < .001, 95% *CI* [-0.819, -0.324]), and day 1 and day 4 (*M* = 5.43, *SE* = 0.085; β = 0.570, *SE* = 0.092, *t*(120) = 6.17, *p* < .001, 95% *CI* [-0.817, -0.321]). No significant differences were found between later days (day 2 vs. day 3, day 2 vs. day 4, day 3 vs. day 4; all *ps* > .999). In contrast, no significant changes in *d′* were found across days in the pair group (day 1: *M* = 5.41, *SE* = 0.081; day 2: *M* = 5.42, *SE* = 0.081; day 3: *M* = 5.45, *SE* = 0.081; day 4: *M* = 5.48, *SE* = 0.081; all *ps* > .999).

Pairwise comparisons between the groups were conducted for each day. On day 1, participants in the pair group showed significantly higher *d′* than those in the individual group (β = 0.550, *SE* = 0.119, *t*(104) = 4.61, *p* < .001, 95% *CI* [-0.785, -0.313]). However, no significant group differences were observed on subsequent days (all *ps* > .660).

#### Response Time

A linear mixed-effects model (LMM) was applied with a Type III Wald χ² test. The model included group (individual vs. pair) and day (1, 2, 3, 4) as fixed effects. Participant was included as a random effect with by-participant random slopes for day. The analysis revealed a significant main effect of day (Wald χ²(3) = 9.46, *p* = .024), while the main effect of group (Wald χ²(1) = 1.90, *p* = .169) and the group × day interaction (Wald χ²(3) = 0.35, *p* = .950) were not significant. Bayesian model comparison strongly supported the inclusion of the main effect of day (*BF*_10_ > 1000). In contrast, the group × day interaction (*BF*_10_ < 0.001) and the main effect of group (*BF*_10_ = 0.028) were not supported, indicating evidence for excluding these effects.

Pairwise comparisons revealed significant decrease in response time across days when averaging over both groups. Compared to day 1 (*M* = 576 ms, *SE* = 10), response time was significantly shorter on day 2 (*M* = 559 ms, *SE* = 9; β = 0.017, *SE* = 0.005, *t*(39.1) = 3.17, *p* = .018, 95% *CI* [0.002, 0.032]), day 3 (*M* = 548 ms, *SE* = 8; β = 0.028, *SE* = 0.007, *t*(40.1) = 3.86, *p* = .002, 95% *CI* [0.008, 0.049]), and day 4 (*M* = 543 ms, *SE* = 9; β = 0.034, *SE* = 0.008, *t*(40.2) = 4.20, *p* < .001, 95% *CI* [0.012, 0.056]). No significant differences were found between day 2 and day 3, day 2 and day 4, or day 3 and day 4 (all *ps* > .084).

### Identification task

#### Accuracy

To examine accuracy in the identification task during learning sessions, a generalized linear mixed model (GLMM) with a binomial link function was fitted, and a Type III Wald χ² test was used to evaluate the effects. The model included group (individual vs. pair), stimulus (rising vs. falling), and day (1, 2, 3, 4) as fixed effects. Participant was included as a random effect with a random intercept. The analysis revealed a significant main effect of stimulus (Wald χ²(1) = 5.10, *p* = .024). Participants exhibited a higher accuracy for sounds with a rising tone (*M* = 97.1%, *SE* = 0.76) than sounds with a falling tone (*M* = 96.4%, *SE* = 0.93). A main effect of day was also observed (Wald χ²(3) = 127.99, *p* < .001) and was modulated by a significant day × group interaction (Wald χ²(3) = 14.65, *p* = .002). No other significant main or interaction effects were found (all *ps* > .085). Bayesian model comparison strongly supported the inclusion of the main effect of day (*BF*_10_ > 1000). In contrast, the interaction between day and group (*BF*_10_ = 0.783) was not supported. The main effect of group (*BF*_10_ = 0.008), the group × stimulus × day interaction (*BF*_10_ < 0.001), and the stimulus × day interaction (*BF*_10_ < 0.001) all showed evidence against inclusion.

Pairwise comparisons were conducted to examine the main effect of day. Response averaged over group and stimulus showed that accuracy differed between day 1 (*M* = 92.5%, *SE* = 0.02) and day 2 (*M* = 96.7%, *SE* = 0.01; β = –0.882, *SE* = 0.072, *z* = –12.17, *p* < .001, 95% *CI* [–1.073, –0.690]), between day 1 and day 3 (*M* = 97.7%, *SE* = 0.01; β = –1.225, *SE* = 0.078, *z* = –15.74, *p* < .001, 95% *CI* [–1.430, –1.019]), and between day 1 and day 4 (*M* = 98.2%, *SE* = 0.01; β = –1.464, *SE* = 0.083, *z* = –17.58, *p* < .001, 95% *CI* [–1.684, –1.244]). Accuracy also differed between day 2 and day 3 (β = –0.343, *SE* = 0.084, *z* = –4.09, *p* < .001, 95% *CI* [–0.565, –0.122]), and between day 2 and day 4 (β = –0.582, *SE* = 0.089, *z* = –6.54, *p* < .001, 95% *CI* [– 0.817, –0.348]). The difference between day 3 and day 4 did not reach significance (*p* = .062).

#### Response Time

Since response times in the identification task were normally distributed, a linear mixed-effects model (LMM) was applied with a Type III Wald χ² test. The model included group (individual vs. pair), stimulus (rising vs. falling), and day (1, 2, 3, 4) as fixed effects. Participant was included as a random effect, with a by-participant random slope for day. The analysis revealed a significant main effect of stimulus (Wald χ²(1) = 63.20, *p* < .001). Response times were faster for sounds with a rising tone (*M* = 1380 ms, *SE* = 16.6) than for those with a falling tone (*M* = 1420 ms, *SE* = 16.6). A significant main effect of day (Wald χ²(3) = 26.15, *p* < .001) was also observed. No other significant main or interaction effects were found (all *ps* > .160). Bayesian model comparison strongly supported the inclusion of the main effect of day (*BF*_10_ > 1000). In contrast, the day × group interaction (*BF*_10_ = 0.035) was not supported. Likewise, the main effect of group (*BF*_10_ = 0.087), the group × stimulus × day interaction (*BF*_10_ < 0.001), and the stimulus × day interaction (*BF*_10_ = 0.003) all showed evidence against inclusion.

Pairwise comparisons revealed significant decreases in response time across multiple days: day 1 (*M* = 1470 ms, *SE* = 18.2) and day 2 (*M* = 1410 ms, *SE* = 19.6; β = 0.0588, *SE* = 0.013, *t*(34.5) = 4.53, *p* < .001, 95% *CI* [0.022, 0.095]), day 1 and day 3 (*M* = 1370 ms, *SE* = 16.8; β = 0.0928, *SE* = 0.0135, *t*(34.7) = 6.89, *p* < .001, 95% *CI* [0.055, 0.131]), day 1 and day 4 (*M* = 1350 ms, *SE* = 17.0; β = 0.1141, *SE* = 0.0145, *t*(36.5) = 7.85, *p* < .001, 95% *CI* [0.074, 0.155]), day 2 and day 3 (β = 0.0341, *SE* = 0.0084, *t*(39.9) = 4.06, *p* = .001, 95% *CI* [0.011, 0.057]), and day 2 and day 4 (β = 0.0553, *SE* = 0.0095, *t*(40.0) = 5.80, *p* < .001, 95% *CI* [0.029, 0.082]). The difference between day 3 and day 4 was also significant (β = 0.0213, *SE* = 0.0060, *t*(40.1) = 3.53, *p* = .006, 95% *CI* [0.005, 0.038]).

## Results from the pre- and post-tests

Behavioral task results indicating learning effects are summarized and shown in Figure 3. Grand-averaged waveforms and topographic maps of MMN and P3a responses, which reflect learning effects, are presented in Figure 4. Additionally, Figure 5 illustrates grand-averaged waveforms and topographic maps of MMN in the passive condition, as well as N2b and P3b in the active condition, showing stimulus-related effects.

**Figure 3.**
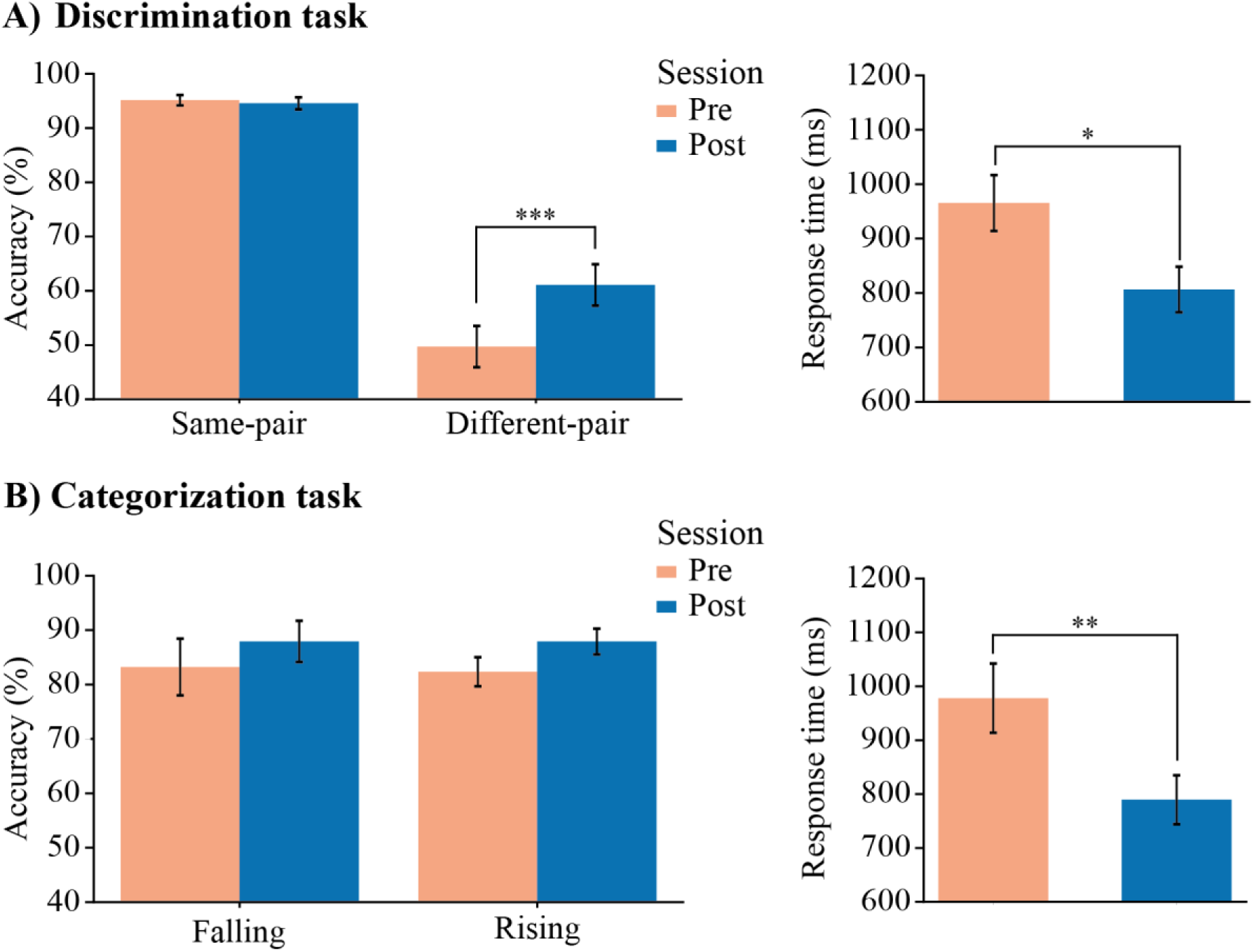
Behavioral results during the pre- and post-tests. A) Results from the discrimination task. Post hoc results for the interaction effect between stimulus and session for accuracy (left) and results for the main effect of session for response time (right) are shown. Group differences were not observed in the analyses. Only significant effects reflecting learning-related changes are marked with asterisks. B) Results from the categorization task. For accuracy (left), no significant differences were observed. For response time (right), statistical results investigating the main effect of time were illustrated. All bar charts display the estimated marginal means (EMMs), with error bars indicating the standard error (*SE*) of the EMMs. * = p < .05, ** = p < .01, *** = p < .001.

**Figure 4.**
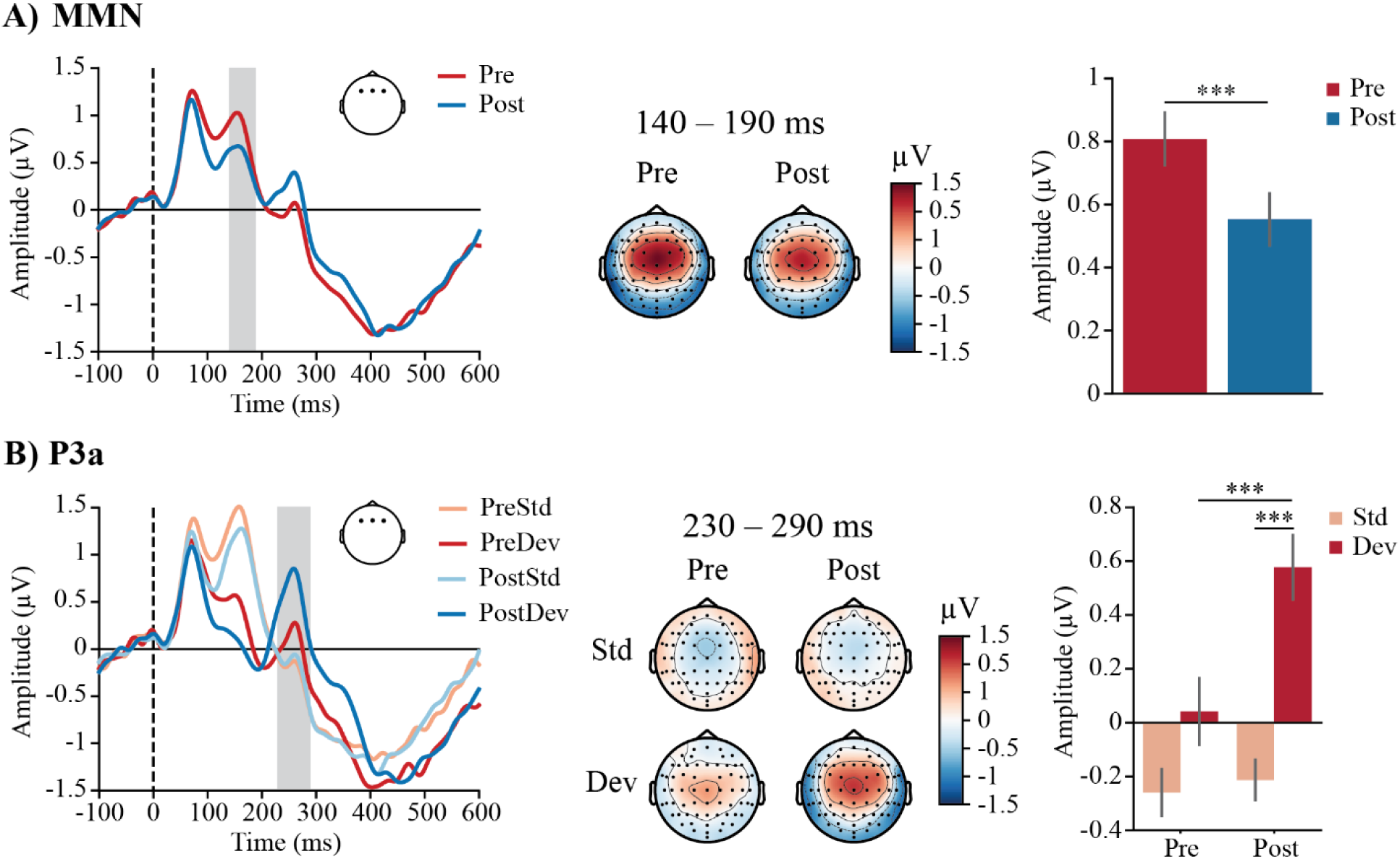
Grand-averaged waveforms, topographic maps, and bar plots of MMN and P3a responses showing learning-related effects. A) Results of MMN investigating the main effect of session. MMN responses are averaged over stimulus types, deviant types, and groups. B) Post hoc results investigating the interaction effect between stimulus type and session for P3a. For A) and B), the grey shaded area in the waveform indicates the time window used for statistical analysis, and the electrode map highlights the electrode pool included in the analysis. In the bar plots, error bars represent the standard error of the mean (SEM). Pre = pre-test, Post = post-test, Std = standard stimuli, Dev = deviant stimuli; *** = p < .001.

**Figure 5.**
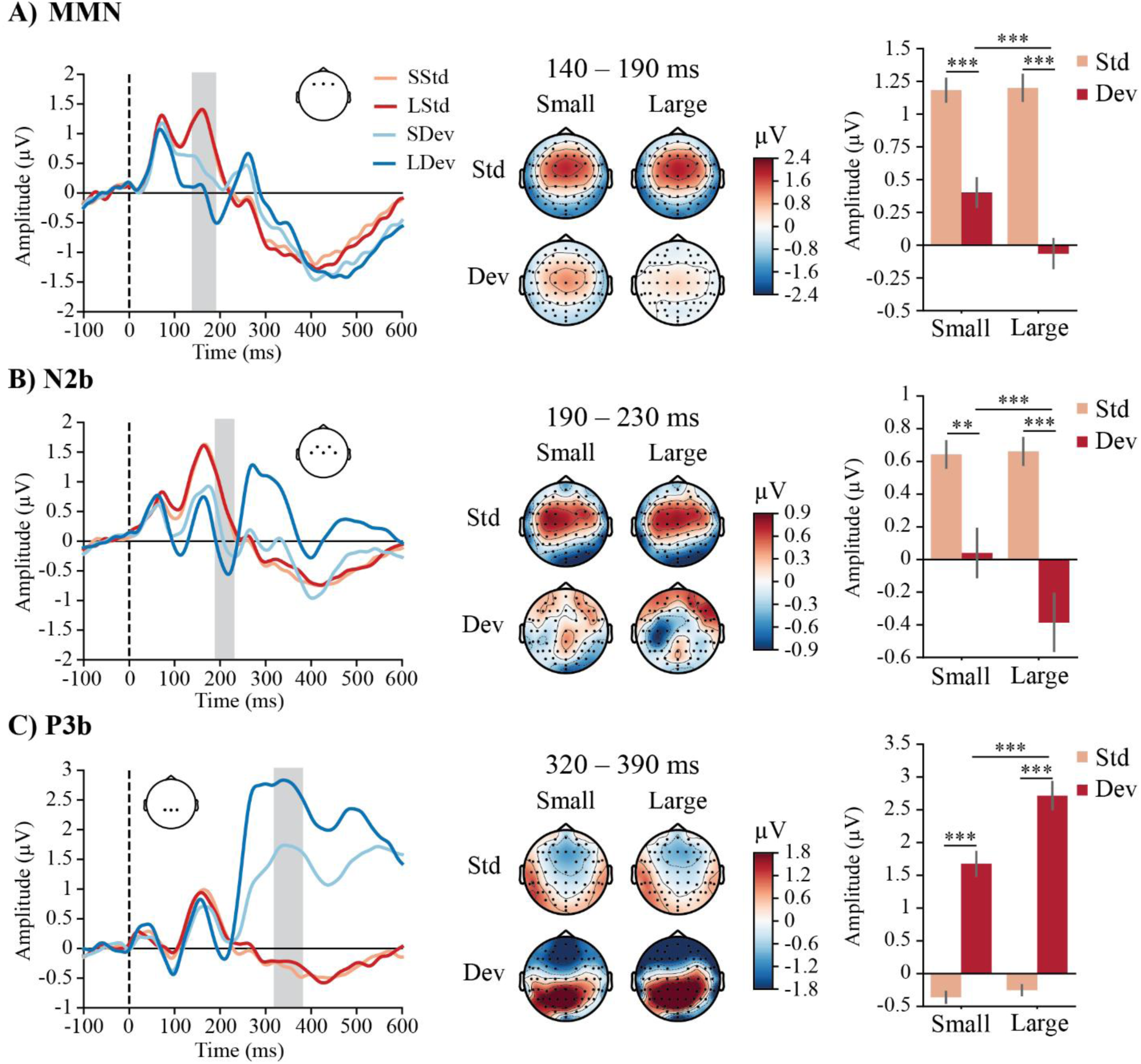
Grand-averaged waveforms and topographic maps of MMN, N2b, and P3b responses, showing stimulus-related effects. The grey shaded area in the waveform indicates the time window used for statistical analysis, and the electrode map highlights the electrode pool included in the analysis. In the bar plots, error bars represent the standard error of the mean (SEM). S = small, L = large, Std = standard stimuli, Dev = deviant stimuli; ** = p < .01, *** = p < .001.

### Behavioral responses in discrimination task

#### Accuracy

To examine accuracy in the discrimination task, a generalized linear mixed model (GLMM) was fitted with group (individual vs. pair), stimulus (same vs. different), and session (pre vs. post) as fixed effects. Participant was included as a random effect, with by-participant random slopes for stimulus and session. The analysis revealed significant main effects of stimulus (Wald χ²(1) = 37.71, *p* < .001) and session (Wald χ²(1) = 14.57, *p* < .001). These main effects were further modified by a significant stimulus × session interaction (Wald χ²(1) = 16.19, *p* < .001). No other significant main or interaction effects were observed (all *ps* > .127). Bayesian model comparison strongly supported the inclusion of the main effect of session (*BF*_10_ > 1000) and the stimulus × session interaction (*BF*_10_ > 1000). In contrast, the main effect of group (*BF*_10_ = 0.011), the group × session interaction (*BF*_10_ = 0.052), and the stimulus × session × group interaction (*BF*_10_ = 0.027) all showed evidence against inclusion.

Pairwise comparisons revealed that accuracy was significantly higher for same-pair trials compared to different-pair trials in both pre-test (β = -2.986, *SE* = 0.233, *z* = -12.797, *p* < .001, 95% *CI* [-3.602, -2.371]) and post-test (β = -2.406, *SE* = 0.232, *z* = -10.357, *p* < .001, 95% *CI* [-3.018, -1.793]). Additionally, the accuracy improvement from pre- to post-test was significant for different-pair trials (*β* = 0.462, *SE* = 0.072, *z* = 6.405, *p* < .001, 95% *CI* [0.272, 0.653]), while no significant change was observed for same-pair trials (*p* > .999). Accuracy for same-pair trials was 95.1% (*SE* = 9.52) at pre-test and 94.6% (*SE* = 11.1) at post-test, whereas for different-pair trials, accuracy was 49.7% (*SE* = 38.1) at pre-test and 61.1% (*SE* = 38.0) at post-test.

#### Response Time

A generalized linear mixed model (GLMM) with a Gamma distribution and log link function was applied to RTs. The model included group (individual vs. pair), stimulus (same vs. different), and session (pre vs. post) as fixed effects, with participant as a random effect, including by-participant random slopes for stimulus and session. The analysis revealed a significant main effect of stimulus, χ²(1) = 6.94, *p* = .008, with participants responding faster to same-pair trials (*M* = 825 ms, *SE* = 39.9) than to different-pair trials (*M* = 939 ms, *SE* = 42.4). A significant main effect of session was also found, χ²(1) = 4.88, *p* = .027, indicating that response times decreased from pre-test (*M* = 964 ms, *SE* = 47.0) to post-test (*M* = 804 ms, *SE* = 38.3). In addition, there was a significant stimulus × session interaction, χ²(1) = 9.46, *p* = .002, and a significant three-way interaction between group, stimulus, and session, χ²(1) = 4.64, *p* = .031. No other main or interaction effects were significant (all *ps* > .763).

Bayesian model comparison strongly supported the inclusion of the main effect of session (*BF*_10_ > 1000). In contrast, the main effect of group (*BF*_10_ = 0.017), the group × session interaction (*BF*_10_ = 0.509), the stimulus × session interaction (*BF*_10_ = 0.054), and the three-way interaction of stimulus × session × group (*BF*_10_ = 0.174) all showed evidence against inclusion.

### Behavioral responses in categorization task

#### Accuracy

To examine accuracy in the categorization task, a generalized linear mixed model (GLMM) with a Type III Wald χ² test was fitted, including group (individual vs. pair), stimulus (rising vs. falling), and session (pre vs. post) as fixed effects. Participant was included as a random effect, with by-participant random slopes for stimulus and session. The analysis revealed a significant stimulus × session interaction (Wald χ²(1) = 5.06, *p* = .024) and a significant group × stimulus × session interaction (Wald χ²(1) = 9.54, *p* = .002). No other significant main or interaction effects were found (all *ps* > .204). Bayesian model comparison supported the inclusion of the main effect of session (*BF*_10_ = 43.051) and provided evidence for the three-way interaction (*BF*_10_ = 1.533). In contrast, the group × session interaction (*BF*_10_ = 0.018), the main effect of group (*BF*_10_ = 0.026), and the stimulus × session interaction (*BF*_10_ = 0.020) all showed evidence against inclusion.

Joint Wald χ² tests were conducted to further investigate the three-way interaction effect. When investigating two-way interaction at different levels of stimulus or session, no significant differences were observed (all *ps* > .059). When examining groups separately, a significant stimulus × session interaction was found for both the individual group (Wald χ²(1) = 5.06, *p* = .025) and the pair group (Wald χ²(1) = 4.24, *p* = .040). No other significant main or interaction effects were observed (all *ps* > .058). Following the joint Wald χ² tests, post hoc pairwise comparisons with Bonferroni correction within each group did not reveal any significant differences (all *ps* > .161).

#### Response Time

For the analysis of response times (RTs), a generalized linear mixed model (GLMM) with a Gamma distribution and log link function and a Wald χ² test were applied. The model included group (individual vs. pair), stimulus (rising vs. falling), and session (pre vs. post) as fixed effects, with participant as a random effect, including by-participant random slopes for stimulus and session. The analysis revealed a significant main effect of session (Wald χ²(1) = 7.49, *p* = .006). Response times were significantly decreased in the post-test (*M* = 789 ms, *SE* = 40.4) compared to the pre-test (*M* = 973 ms, *SE* = 58.0). There were no other main or interaction effects (all *ps* > .129). Bayesian model comparison supported the inclusion of the main effect of session (*BF*_10_ > 1000). In contrast, the main effect of group (*BF*_10_ = 0.033), the group × session interaction (*BF*_10_ = 0.039), and the stimulus × session × group interaction (*BF*_10_ = 0.036) were not supported. BIC-based Bayes factors provided only weak evidence for the stimulus × session interaction (*BF*_10_ = 1.468), and the corresponding frequentist test was not significant (*p* = .460). Thus, there was no reliable evidence that learning differed by stimulus type across sessions.

### ERPs in passive listening

#### MMN (140-190 ms)

The MMN amplitudes were analyzed using a four-way repeated-measures ANOVA with stimulus type (deviant vs. standard), deviant type (small vs. large), and session (pre vs. post) as within-subject factors, and group (individual vs. pair) as a between-subject factor. The analysis revealed a significant main effect of session (*F*(1, 39) = 12.695, *p* < .001, partial η² = .246). Participants exhibited a larger response, shifting towards negativity polarity, in the post-test (*M* = 0.551 μV, *SE* = 0.114) compared to the pre-test (*M* = 0.805 μV, *SE* = 0.124). A main effect of stimulus type (*F*(1, 39) = 109.140, *p* < .001, partial η² = .737) and a main effect deviant type (*F*(1, 39) = 18.521, *p* < .001, partial η² = .322) were also observed. These main effects were further modified by a deviant type × group interaction effect (*F*(1, 39) = 4.483, *p* = .041, partial η² = .103) and a stimulus type × deviant type interaction effect (*F*(1, 39) = 12.872, *p* < .001, partial η² = .248). There were no other main or interaction effects (all *ps* > .102).

Post hoc comparisons were conducted to further investigate the interaction effects. For the deviant type × group interaction, a significant difference was found between small change stimuli (averaged across small deviants and immediately preceding standards; *M* = 0.752 μV, *SE* = 0.164) and large change stimuli (averaged across large deviants and immediately preceding standards; *M* = 0.412 μV, *SE* = 0.170; *p* < .001, 95% *CI* [0.129, 0.551], *d* = 0.37) in the individual group, while the pair group did not show similar difference between the stimulus types. The comparisons between the individual and pair groups did not reveal any statistical significance (all *ps* > .999).

For the stimulus type × deviant type interaction, significant differences were observed between small and large deviants (*p* < .001, 95% *CI* [0.238, 0.705], *d* = 0.51), but not for standards (*p* > .999). Large deviants (*M* = -0.069 μV, *SE* = 0.141) elicited greater amplitudes toward negative polarity compared to small deviants (*M* = 0.402 μV, *SE* = 0.143). Furthermore, for both large and small change stimuli, deviants elicited larger responses compared to the corresponding standards (standard immediately preceding small deviants: *M* = 1.182 μV, *SE* = 0.112; *p* < .001, 95% *CI* [0.456, 1.104], *d* = 0.85; standard preceeding large deviants: *M* = 1.198 μV, *SE* = 0.125; *p* < .001, 95% *CI* [0.943, 1.591], *d* = 1.38).

#### P3a (230-290 ms)

A four-way repeated-measures ANOVA with stimulus type (deviant vs. standard), deviant type (small vs. large), and session (pre vs. post) as within-subject factors, and group (individual vs. pair) as a between-subject factor was conducted for P3a amplitudes. Significant main effects were observed for session (*F*(1, 39) = 11.135, *p* = .002, partial η² = .222) and stimulus type (*F*(1, 39) = 25.040, *p* < .001, partial η² = .391). These main effects were further modified by a significant session × stimulus type interaction (*F*(1, 39) = 14.599, *p* < .001, partial η² = .272).

There were no other main or interaction effects (all *p* > .110).

Post hoc analyses revealed that P3a amplitudes to deviants were significantly larger toward positive polarity in the post-test (*M* = 0.574 μV, *SE* = 0.156) compared to the pre-test (*M* = 0.041 μV, *SE* = 0.67; p < .001, 95% *CI* [-0.833, -0.243], *d* = 0.57). For standard stimuli, there was no significant difference between the two test sessions (*p* > .999). Additionally, comparisons that were conducted within each test session showed that amplitudes to deviants were larger than those to standards in the post-tests (*p* < .001, 95% *CI* [-1.135, -0.446], *d* = 0.84), while no differences were observed in the pre-tests (*p* = .119). Mean amplitudes for standard stimuli were -0.258 μV (*SE* = 0.110) in pre-tests and -0.214 μV (*SE* = 0.098) in post-tests.

### Behavioral results and ERPs in active listening

#### Behavioral results

For accuracy, d-prime (*d’*) was applied and analyzed using a linear mixed-effects model (LMM) with a Type III Wald χ² test. The model included group (individual vs. pair) and session (pre vs. post) as fixed effects. Participant was included as a random effect with a random intercept. The analysis revealed a significant main effect of session, χ²(1) = 27.32, *p* < .001. Bayesian model comparison strongly supported this effect (*BF*_10_ > 1000). In contrast, the group × session interaction (*BF*_10_ = 0.117) and the main effect of group (*BF*_10_ = 0.187) were not supported, providing evidence against their inclusion. Participants showed higher d′ values at post-test (*M* = 4.08, *SE* = 0.169) compared to pre-test (*M* = 3.26, *SE* = 0.169).

For response times, a linear mixed-effects model (LMM) with a Type III Wald χ² test was applied. The model included group (individual vs. pair) and session (pre vs. post) as fixed effects, and participant as a random effect with a random intercept. The analysis showed no significant main or interaction effects (all *ps* > .085). However, Bayesian model comparison provided strong evidence for the inclusion of the main effect of session (*BF*_10_ = 9.948). In contrast, the group × session interaction (*BF*_10_ = 0.507) and the main effect of group (*BF*_10_ = 0.114) were not supported, indicating evidence against their inclusion. Participants responded more quickly at post-test (*M* = 409 ms, *SE* = 7) compared to pre-test (*M* = 437 ms, *SE* = 7).

#### N2b (190-230 ms)

The N2b amplitudes were analyzed using a four-way repeated-measures ANOVA with stimulus type (deviant vs. standard), deviant type (small vs. large), and session (pre vs. post) as within-subject factors, and group (individual vs. pair) as a between-subject factor. Significant main effects were observed for deviant type (*F*(1, 38) = 11.718, *p* = .001, partial η² = .236) and stimulus type (*F*(1, 38) = 25.665, *p* < .001, partial η² = .403). These main effects were further modified by a significant deviant type × stimulus type interaction (*F*(1, 38) = 14.861, *p* < .001, partial η² = .281). No other significant main or interaction effects were found (all *p* > .096).

Post hoc analyses revealed that large deviants elicited greater negativity (*M* = -0.400 μV, *SE* = 0.221) compared to small deviants (*M* = 0.033 μV, *SE* = 0.190; *p* < .001, 95% *CI* [0.201, 0.650], *d* = 0.37). For standard stimuli, there was no significant difference between standards before small and standards before large deviants (*p* > .999). Additionally, comparisons between deviants and standards were conducted for large and small stimuli separately. For large stimuli, responses to deviants were more negative than those to standards (*M* = 0.655 μV, *SE* = 0.101; *p* < .001, 95% *CI* [0.571, 1.522], *d* = 0.91). Similarly, for small stimuli, responses to deviants showed greater negativity than those to standards (*M* = 0.640 μV, *SE* = 0.100; *p* = .006, 95% *CI* [0.127, 1.078], *d* = 0.52).

#### P3b (320-390 ms)

The P3b amplitudes were analyzed using a four-way repeated-measures ANOVA with stimulus type (deviant vs. standard), deviant type (small vs. large), and session (pre vs. post) as within-subject factors, and group (individual vs. pair) as a between-subject factor. Significant main effects were observed for deviant type (*F*(1, 38) = 49.202, *p* < .001, partial η² = .564) and stimulus type (*F*(1, 38) = 114.756, *p* < .001, partial η² = .751). These main effects were further modified by a significant deviant type × stimulus type interaction (*F*(1, 38) = 41.169, *p* < .001, partial η² = .520). There were no other main or interaction effects (all *p* > .085).

Post hoc analyses revealed that large deviants elicited significantly more positive amplitudes (*M* = 2.704 μV, *SE* = 0.259) compared to small deviants (*M* = 1.665 μV, *SE* = 0.227; *p* < .001, 95% *CI* [-1.333, -0.741], d = 0.78). No significant difference was observed between the standards (*p* > .999). Additionally, comparisons between deviants and standards were conducted for large and small stimuli separately. For large stimuli, deviants elicited larger responses than standards (*M* = -0.253 μV, *SE* = 0.089; *p* < .001, 95% *CI* [-3.642, -2.293], *d* = 2.24), whereas for small stimuli, responses to deviants were more positive in amplitude than those to standards (*M* = - 0.361 μV, *SE* = 0.103; *p* < .001, 95% *CI* [-2.713, -1.363], *d* = 1.54).

## DISCUSSION

The present study examined how short-term training shapes the perception and neural change detection of foreign Mandarin tones in Finnish adult learners, and whether training in pairs enhances learning efficiency compared to training alone. Consistent with our hypotheses, we observed rapid and robust learning-related changes in both behavioral performance and electrophysiological responses, with ERP effects emerging specifically in passive change detection. In addition, learners who trained in pairs showed an early advantage on the first training day in the change detection task, suggesting that social presence may facilitate the initial stages of phonetic learning. Next, we discuss these findings in detail.

### Behavioral evidence of learning

During the learning phase, participants showed clear improvements in both the change detection and identification tasks, reflected in higher accuracy and faster responses, indicating rapid perceptual attunement to non-native lexical tone contrasts.

Change detection was measured using an oddball condition, in which the tone in the syllable /fa/ occasionally changed, and participants were asked to press a button whenever a change occurred, without indicating the type of change. A group effect was observed for d′: accuracy increased from the first to the second day specifically for individual learners, whereas pair learners exhibited stable, high sensitivity that was already greater than that of individual learners on the first day. No further gains were observed after the second day in the individual group, suggesting that perceptual sensitivity improved substantially during the initial exposure and quickly reached a plateau. Response times in both groups were longest on the first day and decreased on all subsequent days compared to Day 1.

The change detection-related group difference on the first learning day may reflect an advantage of social presence. In the pair group, learners sat next to each other and could see each other’s accuracy scores on the screen. Although direct interaction was limited, this co-presence and shared performance feedback—possibly including a subtle competitive element— may have enhanced engagement and attention (Verga & Kotz, 2013). The benefits of social interaction observed here are consistent with prior findings from vocabulary learning, where interactive conditions, such as playing a word-learning game with a knowledgeable partner, led to faster responses and higher accuracy compared to non-interactive conditions (Verga et al., 2015; Verga & Kotz, 2017, 2019). However, in the present study, this advantage was not maintained across later learning days, as both groups ultimately reached similarly high sensitivity. The limited nature of interaction in the pair group may partly explain why the observed benefit was not more pronounced or sustained. Alternatively, the effect of social interaction may have been less detectable because sensitivity to tonal changes was already high, suggesting a potential ceiling effect or that the task itself was relatively easy, as reflected in the high d′ values.

In the identification task, accuracy improved steadily from the first to the third learning day, with no further improvement from day 3 to day 4, suggesting that most of the learning gains were achieved within the first three sessions. No group differences were found in any of the days. The high accuracy in the identification task indicates that, despite the variability of syllables, tone recognition was relatively easy for the participants. Response times decreased steadily across all days after the initial session and no group differences in them were found. Thus, the predicted faster improvement in change detection relative to identification was evident in accuracy, whereas response times revealed a similar gradual pattern across both tasks.

Perceptual improvements were also evident in the behavioral pre- to post-tests. In the discrimination task, accuracy for different-pair trials increased significantly from around 50% (chance level) to 60%, whereas accuracy for same-pair trials was already high in the pre-test (≈95%) and remained stable after training. The high baseline performance for same pairs likely reflects a response bias toward the “same” option, while the initially low accuracy for different pairs underscores the difficulty of detecting subtle pitch contour differences without established tonal categories. Such asymmetry is well documented among non-tonal language speakers, who often default to a “same” response when uncertain, yielding high same-pair accuracy but lower different-pair accuracy (Francis et al., 2008). Response times decreased for both same- and different-pair trials from pre- to post-test, indicating more efficient perceptual processing after training.

In the categorization task during pre- and post-tests, only response times improved, while accuracy remained relatively high from the pre- to post-test (≈80%). It is notable that in the pre- and post-tests, the categorization task used only the vowel /a/ presented with several different tones, including relatively flat tones, whereas the identification tasks during learning involved multiple syllables with clearly rising and falling tones. In addition, the categorization task began with examples of rising and falling tones, which likely served as perceptual cues. Even if listeners could not detect all subtle pitch details, they could match subsequent stimuli to these cues using broad pitch movement patterns familiar from their native language intonation. In the discrimination task, direct acoustic comparison between two tones without such categorical cues was required, making subtle differences harder to detect and resulting in lower accuracy for different-pair trials.

In the active change detection task during pre- and post-tests, behavioral responses were recorded together with ERPs. Accuracy (d′) increased and response time decreased from pre-test to post-test, with no group differences at either time point. Because pre- and post-tests were conducted individually for all participants, the observed benefit of the pair group in behavioral change detection may have been limited to the learning sessions, when participants were together, supporting the notion that attentional mechanisms underlie this effect.

### Neural correlates of learning

In the passive oddball condition, larger P3a amplitudes to deviants were observed in the post-test compared to those in the pre-test, indicating increased automatic attentional engagement with tonal changes after learning. This enhancement was specific to deviants, with no amplitude change for standards, suggesting that the effect reflected improved processing of tonal changes rather than a general changes in neural responsiveness. These results align with previous findings showing P3a sensitivity to auditory novelty and its amplification due to passive exposure to Chinese tones (Kurkela et al., 2019; Li et al., 2025). Instead, the MMN amplitude showed a general shift toward negative polarity from pre- to post-test. This main effect of session may indicate increased neural responsiveness to acoustic changes, but given the absence of other main effects or interactions, it reflects an overall trend rather than effects tied to specific groups, stimulus types, or deviant types, and should therefore be interpreted with caution.

In the active change detection task, larger N2b and P3b responses were elicited by large deviants than by small deviants, and by deviants than by standards, regardless of test session. No learning-related changes were found in N2b or P3b amplitudes. This differs from some studies reporting P3b enhancement after training or exposure (Alain et al., 2010; Giroud et al., 2017; Kurkela et al., 2019) and from findings that N2b enhancement can follow intensive identification training (Alain et al., 2010). However, other studies have reported decreases or no change in N2b and/or P3b (e.g., Ben-David et al., 2011; Giroud et al., 2017; Kurkela et al., 2019), indicating that learning-related changes in these components can vary depending on factors such as task design and training duration. The level of attentional engagement may also modulate these responses. Yang et al. (2024) found no group differences in N2b and P3b between Finnish and Chinese native speakers in an active change detection task, suggesting that the mechanisms underlying change detection and attentional shifting operate effectively for both native and non-native speech sounds when attention is directed to the stimuli. It is therefore possible that in the present study, participants’ attentive listening during the active task resulted in robust N2b and P3b responses in the pre-test, leaving little room for observable changes after learning.

### Limitations

The present study has several limitations that should be considered. First, there was no control group that completed the tasks without training, so improvements in behavioral responses at whole group level could partly reflect increased familiarity with the tasks. However, the increase in P3a amplitudes, an index of automatic attentional engagement measured during passive listening, is less likely to be influenced by task repetition effects, providing further evidence that participants experienced genuine learning-related improvement with enhanced sensitivity to non-native phonetic contrasts. Due to the relatively small sample size, it was not possible to examine correlations between behavioral responses and neural changes in the present study. Future studies may build on this combined approach to capture both early neural adaptations and how they relate to behavioral improvements. Second, the social interaction in the pair group was limited to co-presence and shared performance feedback, without extensive verbal or collaborative engagement, which may explain the limited group findings. Finally, the sample consisted of Finnish-speaking adult learners with no prior exposure to tonal languages, so the generalizability of the findings to other populations, such as children or speakers of other languages, remains to be tested.

## Conclusion

Our study demonstrates that a four-hour phonetic training protocol enhances adult learners’ sensitivity to non-native lexical tone contrasts, as reflected in behavioral improvements and selective neural changes (P3a enhancement to novel speech sounds) during passive change detection. Social presence provided an early advantage in behavioral change detection during training. These findings underscore the value of short, targeted training and multimodal assessment for capturing multiple dimensions of adult phonetic learning, including transfer to novel speech sounds.

## Data Availability Statement

The research data will be stored in an open repository once the data has been anonymized. The data will be anonymized once the manuscript (this manuscript or a revised version of it) has been authorized for publication.

## Author Contributions

**Xueqiao Li:** Conceptualization (Supporting), Data curation (Lead), Formal analysis (Lead), Funding acquisition (Supporting), Investigation (Lead), Methodology (Equal), Supervision (Equal), Validation (Equal), Visualization (Lead), Writing - original draft (Lead), Writing – review & editing (Equal). **Hyeonjeong Jeong**: Conceptualization (Supporting), Funding acquisition (Supporting), Writing - review & editing (Equal). **Piia Astikainen**: Conceptualization (Lead), Funding acquisition (Lead), Investigation (Supporting), Project administration (Lead), Resources (Lead), Supervision (Equal), Validation (Equal), Writing - original draft (Supporting), Writing - review & editing (Equal).

## Acknowledgments

The authors thank the research assistants and students (Matias Siltakoski, Petrus Kuusisto, Peppi Salminen, Marika Alatalo, Emilia Kokko, Elias Rantakokko, Sohvi Saarinen, Tuomo Perttula, Maria Lampilahti, Titta Holopainen, and Kyösti Sorsa) for their help during data collection, Dr. Viki-Veikko Elomaa and Petri Kinnunen for their technical assistance; and Joona Muotka for his statistical advice.

