## supplementary materials for "Rapid phonetic learning of Mandarin tones in adults: Daily behavioral improvement and brain activity changes"

### Daily state questionnaire during learning sessions: method, results, and summary

#### Methods

During each learning session, participants completed a short questionnaire in Finnish. Two items measured participants' current state before the learning session, and four items were presented after the session to evaluate their state during the learning session, as well as their self-evaluation of how well they learned and how pleasant they felt during learning. All items were rated on a 5-point Likert scale. The details of each item are provided in Table 1.

Table 1. Items of the daily state questionnaire administered before and after each learning session. The original items were in Finnish; only English versions are shown here. The full Finnish version is available upon request.

| Timing | Question | Scale |
|---|---|---|
| **Pre-learning** | Please rate your current alertness level. | Extremely tired – Extremely alert |
| | How motivated are you for the upcoming learning session? | Low motivation – High motivation |
| **Post-learning** | Please rate your current alertness level. | Extremely tired – Extremely alert |
| | Please rate your motivation during the learning session. | Low motivation – High motivation |
| | Please rate how well you learned. | Not at all – Very well |
| | How pleasant was the learning session? | Very unpleasant – Very pleasant |

Statistical analyses of questionnaire data were conducted using IBM SPSS Statistics (version 30.0.0.0; IBM Corporation, Armonk, NY, USA). Repeated-measures ANOVAs were used, and the details of the model structures are reported in the following Results section.

### Results

#### *Alertness*

A three-way repeated-measures ANOVA was conducted with day (1 vs. 2 vs. 3 vs. 4) and session (pre vs. during) as within-subject factors and group (individual vs. pair) as a between-subject factor. A significant main effect of day was found, $F(3, 120) = 3.97$, $p = .010$, partial $\eta^2 = .090$. Post hoc comparisons with Bonferroni correction indicated that participants exhibited significantly higher alertness on Day 3 ($M = 3.50$, SE = 0.098) compared to Day 1 ($M = 3.04$, SE = 0.102, $p = .009$). No significant differences were observed between the other days (all $p > .410$). A significant main effect of session was also found, $F(1, 40) = 79.41$, $p < .001$, partial $\eta^2 = .665$. Participants reported significantly higher alertness before the start of learning sessions ($M = 3.62$, SE = 0.076) than during learning sessions ($M = 2.91$, SE = 0.087) across all days. There were no other main or interaction effects (all $p > .105$). The results are shown in Figure 1.

#### *Motivation*

A similar three-way repeated-measures ANOVA was conducted for participants' self-evaluated motivation. A significant main effect of day was found, $F(3, 120) = 3.21$, $p = .035$, partial $\eta^2 = .074$; however, post hoc comparisons with Bonferroni correction did not reveal any significant differences (all $p > .280$). A significant main effect of session was found, $F(1, 40) = 12.00$, $p = .001$, partial $\eta^2 = .231$, indicating that participants reported higher motivation before the start of learning sessions ($M = 3.72$, SE = 0.108) than during learning sessions ($M = 3.45$, SE = 0.119). No other main or interaction effects were found (all $p > .072$). The results are presented in Figure 1.

#### *Self-assessment of learning*

A two-way repeated-measures ANOVA with day (1 vs. 2 vs. 3 vs. 4) as a within-subject factor and group (individual vs. pair) as a between-subject factor was conducted for participants' self-assessment of learning. A significant main effect of day was found, $F(3, 120) = 3.79$, $p = .012$, partial $\eta^2 = .087$. This effect was modulated by a significant interaction effect of day and group, $F(3, 120) = 3.94$, $p = .010$, partial $\eta^2 = .090$. Post hoc simple effects analysis with Bonferroni correction indicated that participants in the pair group reported significantly higher self-assessments of learning ($M = 3.82$, SE = 0.178) than those in the individual group ($M = 3.15$, SE = 0.186) on Day 2 ($p = .034$). Within-group comparisons revealed that only participants in the Pair group showed significant differences in learning self-assessments across days (Day 1 < Day 2, $p = .034$; Day 2 > Day 4, $p < .001$; Day 3 > Day 4, $p = .032$). No significant differences between days were observed for participants in the individual group (all $p > .999$). There was no main effect of group ($p = .245$). The results are presented in Figure 1.

*Pleasantness During Learning*

A similar two-way repeated-measures ANOVA was conducted for participants' pleasantness during learning. A significant main effect of day was found, $F(3, 120) = 4.10$, $p = .014$, partial $\eta^2 = .093$; however, post hoc comparisons with Bonferroni correction did not reveal any significant differences (all $p > .073$). No other main or interaction effects were found (all $p > .070$).

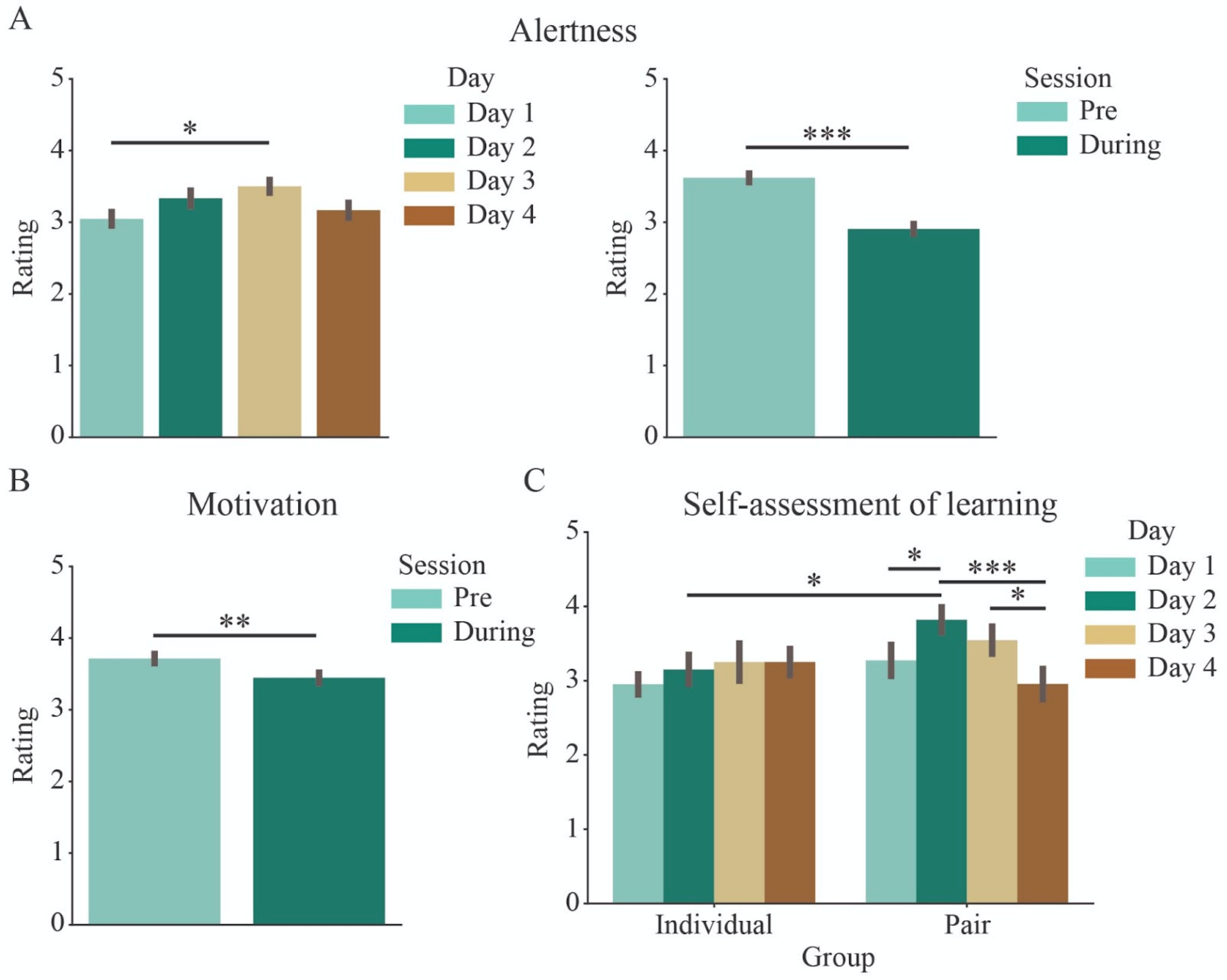


Figure 1. Self-evaluated states during the learning phase. ***$p < .001$, **$p < .01$, *$p < .05$

### Summary

Questionnaire responses indicated some variation in participants' perceived alertness, motivation, self-assessed learning, and pleasantness across the four learning days. For alertness and motivation, participants reported higher ratings before than during learning sessions, possibly reflecting greater energy at the start of each session compared to during the learning activities. Self-assessment of learning showed an interaction between day and group, with participants in the pair group reporting more favorable learning outcomes on Day 2 and some fluctuations across days. These findings suggest changes in perceived state and experience during the learning period, with some indication that learning in pairs may be associated with more positive self-evaluations.

### List of auditory stimuli used in the learning phase

The list of syllables used in the exposure to speech sounds with visual guidance, the identification task, and the listen-and-repeat task is presented in Table 2.

Table 2. List of consonant-vowel syllables used in the learning phase. The syllables were presented in three different tones (flat, rising, falling).

| Consonant–vowel syllables | | | | | | | | | |
|---|---|---|---|---|---|---|---|---|---|
| bai | ban | bian | bin | bo | bu | da | dai | dan | dao |
| de | di | dian | dong | du | gai | gao | ge | gei | gu |
| guang | ha | han | hao | hen | hong | hu | hua | huan | huang |
| kai | kao | ken | kou | ku | kuai | kuan | ma | mai | man |
| men | mi | miao | mo | mu | pai | pao | pi | piao | pu |
| se | si | suan | te | ti | tian | tiao | tong | tou | yan |
